# Modeling magnesium and calcium transport along male rat kidney and the effects of diuretics

**DOI:** 10.1101/2023.10.07.561340

**Authors:** Pritha Dutta, Anita T. Layton

**Author notes:** Correspondence: Pritha Dutta, Department of Applied Mathematics, University of Waterloo Waterloo, Ontario, Canada N2L 3G1.

## Abstract

Calcium (Ca^2+^) and magnesium (Mg^2+^) are essential for cellular function. The kidneys play an important role in maintaining the homeostasis of these cations. Their reabsorption along the nephron is dependent on distinct trans- and paracellular pathways, and is coupled to the transport of other electrolytes. Notably, sodium (Na^+^) transport establishes an electrochemical gradient to drive Ca^2+^ reabsorption. Consequently, alterations in renal Na^+^ handling, under pathophysiological conditions or pharmacological manipulations, can have major effects on Ca^2+^ transport. One such condition is the administration of diuretics, which are used to treat a large range of clinical conditions, but most commonly for the management of blood pressure and fluid balance. While the pharmacological targets of diuretics typically directly mediate Na^+^ transport, they also indirectly affect renal Ca^2+^ and Mg^2+^ handling, i.e., by establishing a prerequisite electrochemical gradient. Thus, substantial alterations in divalent cation handling can be expected following diuretic treatment. To investigate renal Ca^2+^ and Mg^2^ handling, and how those processes are affected by diuretics treatment, we have developed sex-specific computational models of electrolyte transport along the nephrons. Model simulations indicate that along the proximal tubule and thick ascending limb, the transport of Ca^2+^ and Mg^2+^ occusr in parallel with Na^+^, but those processes are dissociated along the distal convoluted tubule. We also simulated the effects of acute administration of loop, thiazide, and K-sparing diuretics. The model predicted significantly increased Mg^2+^ excretion, no significant alteration in Mg^2+^ excretion, and significantly decreased Mg^2+^ excretion on treatment with loop, thiazide, and K-sparing diuretics, respectively, in agreement with experimental studies. The present models can be used to conduct *in silico* studies on how the kidney adapts to alterations in Ca^2+^ and Mg^2+^ homeostasis during various physiological and pathophysiological conditions, such as pregnancy, diabetes, and chronic kidney disease.

## 1. Introduction

The divalent cations, Ca^2+^ and Mg^2+^, are important for various physiological processes. About 99% of the body Ca^2+^ is stored in bones, where it forms a calcium-phosphate compound called hydroxyapatite [1]. The remaining 1% of body calcium plays an important role in various other physiological processes such as cell signaling, both skeletal and smooth muscle contraction, and blood clotting [1]. Mg^2+^ plays a pivotal role in energy-demanding metabolic reactions, protein synthesis, ensuring membrane integrity, facilitating nervous tissue conduction, promoting neuromuscular excitability, regulating muscle contraction, influencing hormone secretion, and participating in intermediary metabolism. Nearly 99% of the body’s Mg^2+^ is distributed within cells or stored in bone, with only a small fraction in circulation [2]. Tight regulation of the serum Ca^2+^ and Mg^2+^ concentrations is essential since too much or too little Ca^2+^ or Mg^2+^ can have dangerous, potentially fatal consequences. To maintain Ca^2+^ balance, it is crucial to regulate the fluxes of Ca^2+^ among the primary organs involved in calcium regulation, namely the intestine, bone, and kidneys. Similarly, both the small intestine and kidneys increasing their fractional Mg^2+^ absorption when magnesium deficiency is detected. Should Mg^2+^ depletion persist, the body’s bone stores step in to help maintain serum magnesium levels by exchanging a portion of their content with extracellular fluid.

The majority, ∼60-70% of the filtered Ca^2+^, is reabsorbed along the proximal tubule through the paracellular pathway [3]. By contrast, paracellular Mg^2+^ permeability in the proximal tubule is very low and hence only 10-25% of the filtered Mg^2+^ is reabsorbed along this segment [3]. The majority of the filtered Mg^2+^ is reabsorbed along the cortical thick ascending limb (60-70%) paracellularly [3]. The paracellular fractional reabsorption of Ca^2+^ along the thick ascending limb is ∼15-25% [3]. The distal convoluted tubule in the final segment that reabsorbs Mg^2+^; hence it plays and important role in fine-tuning urinary Mg^2+^ excretion. 5-10% of the filtered Mg^2+^ is reabsorbed transcellularly along the distal convoluted tubule mediated by the transient receptor potential melastatin 6 (TRPM6) on the apical membrane and Na^+^/Mg^2+^ exchanger on the basolateral membrane [3]. 5-10% of the filtered Ca^2+^ is reabsorbed transcellularly along the distal convoluted tubule and connecting tubule mediated by the transient receptor potential vanilloid 5 (TRPV5) on the apical membrane, and Na^+^/Ca^2+^ exchanger (NCX1) and plasma membrane Ca^2+^-ATPase (PMCA) on the basolateral membrane [3]. Finally, 2-5% of the filtered Mg^2+^ and Ca^2+^ are excreted through urine [3].

Our understanding of Ca^2+^ and Mg^2+^ handling within different segments of the nephron has been greatly advanced through micropuncture and microperfusion studies in rodent nephrons [4,5]. Furthermore, recent genetic studies has expanded our knowledge about the protein mediators of Ca^2+^ and Mg^2+^ transport [6]. Despite these advances, our understanding of renal handling of these electrolytes remains incomplete. What fraction of the renal reabsorption goes through the transcellular versus paracellular pathway? To what extent is the renal transport in each nephron segment coupled to the transport of other electrolytes, e.g., Na^+^, K^+^, and Cl^-^? To answer these questions, we developed a detailed computational model epithelial transport of electrolytes and water along the nephrons in a rat kidney, we and conducted simulations to predict the renal transport of Ca^2+^ and Mg^2+^ as well as other electrolytes and water under different physiological conditions.

Besides electrolyte and fluid homeostasis, the kidney also plays an essential role in maintaining a normal blood pressure. For the management of blood pressure and fluid balance, diuretics are commonly prescribed. Although the pharmacological targets of diuretics directly affect Na^+^ transport, they also indirectly affect renal Mg^2+^ and Ca^2+^ reabsorption through changes in electrochemical gradient. How is renal Ca^2+^ and Mg^2+^ transport affected by the administration of diuretics? To answer this question, we simulate the effect of acute administration of three classes of diuretics-loop, thiazide, and K-sparing diuretics, on renal Mg^2+^ and Ca^2+^ transport and excretion.

## 2. Methods

We previously developed an epithelial cell-based model of solute transport along different nephron populations of a rat kidney in a male rat [7–10]. The model represents six classes of nephrons: a superficial nephron and five juxtamedullary nephrons. The superficial nephrons account for two-thirds of the nephron population, and they extend from Bowman’s capsule to the papillary tip. The remaining one-third are juxtamedullary nephrons, which possess loops of Henle that reach to different depths in the inner medulla; most of the long loops turn within the upper inner medulla. Baseline single nephron glomerular filtration rate (SNGFR) is set to ∼30 and ∼45 nL/min for the superficial and juxtamedullary nephrons, respectively. The model superficial nephron includes ten functionally distinct segments: proximal convoluted tubule, proximal straight tubule or S3, short descending limb, medullary thick ascending limb, cortical thick ascending limb, distal convoluted tubule, connecting tubule, cortical collecting duct, outer-medullary collecting duct, and inner-medullary collecting duct. Each juxtamedullary nephron includes also the long descending limb and long ascending limb segments. The water and solute transport processes along each nephron segment are simulated by a number of epithelial cell models connected in series. Each cell model includes the apical (lumen-facing) and basolateral (interstitium-facing) membranes, and is separated from the adjacent cell by a paracellular space. A schematic diagram for the model nephrons is shown in Fig. X.

Transport capacity of the juxtamedullary proximal tubule is assumed to be 75% higher than that of the superficial proximal tubule, such that fluid flow into the S3 segment is predicted to be between 11-13 nL/min. The model accounts for the following 17 solutes: Na^+^, K^+^, Cl^-^, HCO_3_^-^, H_2_CO_3_, CO_2_, NH_3_, NH_4_^+^, HPO_4_^2-^, H_2_PO_4_^-^, H^+^, HCO_2_^-^, H_2_CO_2_, urea, glucose, Ca^2+^, and, in this study, Mg^2+^ as well. The model is formulated for the steady state and predicts luminal fluid flow, hydrostatic pressure, luminal fluid solute concentrations, and, with the exception of the descending limb segment, cytosolic solute concentrations, membrane potential, transcellular and paracellular fluxes, and urinary excretion.

Below we summarize model representation of Mg^2+^ transport along the proximal tubule, cortical thick ascending limb, and distal convoluted tubule. Model parameters that describe Mg^2+^ transport are given in Table 1. The analogous model description and parameter for Ca^2+^ transport cab be found in Ref. [11]. Additional model parameters can be found in Ref. [12].

**Table 1.** Mg^2+^-specific parameters for all the segments along the superficial nephron. Values marked (*) are adjusted and those marked (**) are the same for both male and female models. PT, proximal tubule; TAL, thick ascending limb; DCT, distal convoluted tubule; CNT, connecting tubule; CD, collecting duct; OMCD, outer-medullary collecting duct; IMCD, inner-medullary collecting duct.

| Parameter | Value |  |
| --- | --- | --- |
|  | Male | Female |
| **PT** |  |  |
| Tight junction permeability to $Mg^{2+}$ at the lumen-LIS interface ($P_{Mg}^{PT,LI}$) | $1.1 \times 10^{-5} \text{ cm/s}$ (**) | $1.1 \times 10^{-5} \text{ cm/s}$ [60] |
| Reflection coefficient of tight junction to $Mg^{2+}$ | 0.89 (*) | 0.89 (**) |
| **cTAL** |  |  |
| Tight junction permeability at the lumen-LIS interface in the absence of $Mg^{2+}$ ($P_{Mg}^{TAL,LI*}$) | $38 \times 10^{-5} \text{ cm/s}$ (*) | $56 \times 10^{-5} \text{ cm/s}$ (*) |
| Maximum half concentration of $Ca^{2+}$ ($EC_{50,Ca}$) | 1.25 mM [25] | 1.25 (**) |
| Maximum half concentration of $Mg^{2+}$ ($EC_{50,Mg}$) | 2.5 mM [25] | 2.5 (**) |
| Hill function coefficient, n | 4 [25] | 4 (**) |
| Inhibitory coefficient of $Ca^{2+}$ on tight junction permeability ($\alpha_{p,Ca}$) | -4/7 [61] | -4/7 (**) |
| Inhibitory coefficient of $Mg^{2+}$ on tight junction permeability ($\alpha_{p,Mg}$) | -0.34 (*) | -0.34 (**) |
| Inhibitory coefficient of $Ca^{2+}$ on NKCC2 activity ($\alpha_{NKCC2,Ca}$) | -0.4 [62] | -0.4 (**) |
| Inhibitory coefficient of $Mg^{2+}$ on NKCC2 activity ($\alpha_{NKCC2,Mg}$) | -0.24 (*) | -0.24 (**) |
| Inhibitory coefficient of $Ca^{2+}$ on ROMK activity ($\alpha_{ROMK,Ca}$) | -0.8 [63,64] | -0.8 (**) |
| Inhibitory coefficient of $Mg^{2+}$ on ROMK activity ($\alpha_{ROMK,Mg}$) | -0.48 (*) | -0.48 (**) |
| **DCT** |  |  |
| TRPM6 channel density ($N_{TRPM6}$) | $22.5 \times 10^4$<br>1/ cm<sup>2</sup> (*) | $45 \times 10^4$ 1/cm<sup>2</sup> (*) |
| Single channel conductance of TRPM6 at pH 7.4 ($g_{pH7.4}$) | 83.6 pS [21] | 83.6 pS (**) |
| Luminal pH for half-maximal conductance of TRPM6 ($pH_{1/2}$) | 4.3 [21] | 4.3 (**) |
| Maximum $Mg^{2+}$ flux through $Na^+/Mg^{2+}$ exchanger ($J_{Mg}^{NaMgX,max}$) | $8.6 \times 10^{-9}$ mmol/cm<sup>2</sup>/s (*) | $9 \times 10^{-9}$ mmol/cm<sup>2</sup>/s (*) |
| Intracellular $Mg^{2+}$ half-saturation constant ($K_{M,MgC}$) | 3.59 M [20,22] | 3.59 M (**) |
| Extracellular $Mg^{2+}$ half-saturation constant ($K_{M,MgS}$) | 1.3 mM [20,22] | 1.3 mM (**) |
| Intracellular $Na^+$ half-saturation constant ($K_{M,NaC}$) | 12.29 mM [20,22] | 12.29 mM (**) |
| Extracellular $Na^+$ half-saturation constant ($K_{M,NaS}$) | 87.5 mM [20,22] | 87.5 mM (**) |
| Excitatory coefficient of $Ca^{2+}$ on NCC activity ($\alpha_{NCC,Ca}$) | 0.5 [65] | 0.5 (**) |
| Excitatory coefficient of $Mg^{2+}$ on NCC activity ($\alpha_{NCC,Mg}$) | 0.3 (*) | 0.3 (**) |
| **CD** |  |  |
| Ca<sup>2+</sup> promoting coefficient for apical<br>HATPase activity of type A OMCD cells<br>($\alpha_{\text{HATP,Ca}}$) | 2 [61] | 2 (**) |
| Mg<sup>2+</sup> promoting coefficient for apical<br>HATPase activity of type A OMCD cells<br>($\alpha_{\text{HATP,Mg}}$) | 1.2 (*) | 1.2 (**) |
| Ca<sup>2+</sup> inhibitory coefficient for apical<br>water permeability of IMCD cells ($\alpha_{\text{Pf,Ca}}$) | -3/8 [61] | -3/8 (**) |
| Mg<sup>2+</sup> inhibitory coefficient for apical<br>water permeability of IMCD cells ($\alpha_{\text{Pf,Mg}}$) | -0.225 (*) | -0.225 (**) |

### 2.1 Mg^2+^ transport along the proximal tubule

The proximal tubule reabsorbs 15-25% of the filtered Mg^2+^ load through the paracellular pathway, which is mediated by claudin-2 and -12 [13] and is driven by the favorable electrochemical gradient established by Na^+^/H^+^ exchanger 3 (NHE3)-mediated Na^+^ reabsorption [5,14,15]. Paracellular electro-diffusive Mg^2+^ flux 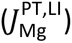 is given by

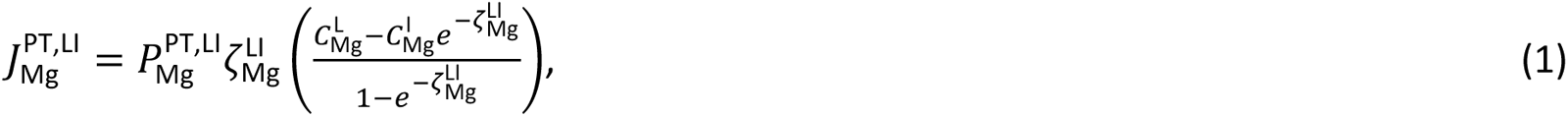

where the superscripts L and I denote lumen and lateral intercellular space (LIS), respectively; 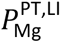 denotes the permeability of Mg^2+^ at the lumen and LIS interface; 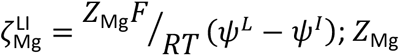 is the valence of Mg^2+^ (+2); 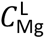 and 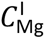 denote Mg^2+^ concentrations in the lumen and LIS, respectively;, ψ*^L^* and ψ*^I^* denote the luminal and LIS membrane potentials, respectively; RT = 2.57 J/mmol; and F = 96.5 C/mmol represents Faraday’s constant.

### 2.2 Mg^2+^ transport along the thick ascending limb

The medullary thick ascending limb has negligible Mg^2+^ reabsorption [16–18]. In contrast, the cortical thick ascending limb reabsorbs 60-70% of the filtered Mg^2+^ via the paracellular route. That flux is mediated by claudins 16 and 19 [19], and is driven by the electrochemical gradient established by Na^+^-K^+^-Cl^-^ cotransporter 2 (NKCC2)-mediated Na^+^ transport [14,15]. Paracellular Mg^2+^ transport in the cortical thick ascending 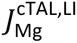 is represented by an expression analogous to Eq (1), with the superscript “PT” replaced by “cTAL”.

### 2.3 Mg^2+^ transport along the distal convoluted tubule

Unlike the proximal tubule and cortical thick ascending limb, where Mg2+ transport proceeds passively via the paracelluar route, the reabsorptive process along the distal convoluted tubule and active and transcellular, and is mediated by the TRPM6 channels, expressed on the apical membranes [20]. Mg^2+^ flux through TRPM6 is given by

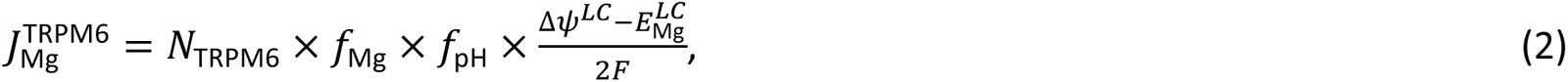

*f*_Mg_ and *f*_pH_ describe the effects of intracellular Mg^2+^ concentration (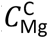) and extracellular pH on TRPM6 [21], and are given by

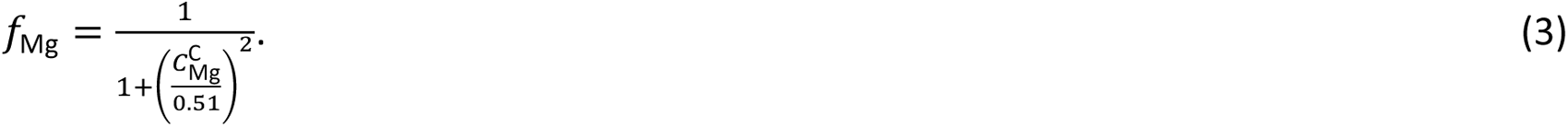

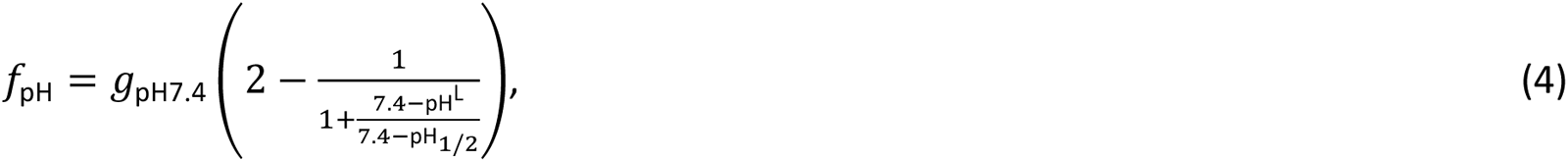

where *g*_pH7.4_ denotes the single channel conductance of TRPM6 at pH 7.4, pH^L^ denotes the luminal fluid pH, and pH_1/2_ denotes the luminal fluid pH for half-maximal conductance. Mg^2+^ efflux through the basolateral membrane is assumed to be mediated an Na^+^/Mg^2+^ exchanger [20,22], given by

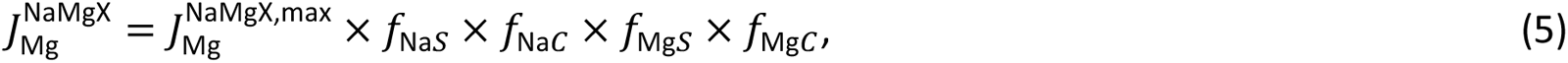

where 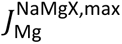 denotes the maximum Mg^2+^ flux through Na^+^/Mg^2+^ exchanger, and the *f* terms represent regulation by extracellular Na^+^, 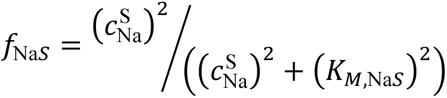, intracellular Na^+^, 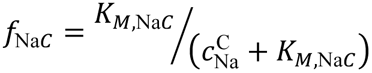, extracellular, Mg^2+^, 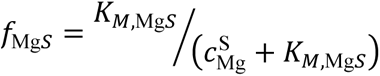, and intracellular Mg^2+^, 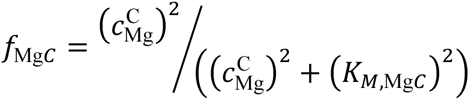. In these expressions, *K_M_*_,Na*S*_, *K_M_*_,Na*C*_, *K_M_*_,Mg*S*_, and *K_M_*_,Mg*C*_ denote the Michaelis-Menten constants.

### 2.4 Calcium-sensing receptor

The calcium-sensing receptor (CaSR) regulates not only Ca^2+^ and Mg^2+^ reabsorption in the kidneys but other electrolytes and water as well by modifying transporter activities. Ca^2+^ is the primary ligand for activating CaSR. At equimolar concentrations, Mg^2+^ is 1/2 to 2/3 as potent as Ca^2+^ in activating CaSR [23,24]. We model the effect of CaSR on a given parameter *v* (*v* may denote paracellular permeability, NKCC2 activity, ROMK activity, or NCC activity; see below) with the following expression:

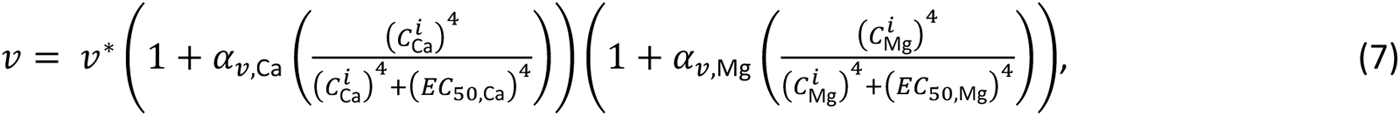

where *v*^∗^ is the value of *v* in the absence of the effect of CaSR, 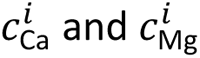 denote the concentration of Ca^2+^ and Mg^2+^ in the luminal (*i*=L) or interstitial (*i*=S) fluid, and *EC*_50,Ca_ = 1.25 mM and *EC*_50,Mg_ = 2.5 mM represent the half-maximal concentrations for Ca^2+^ and Mg^2+^ [25], respectively. CaSR is ubiquitously expressed in the kidney both along the apical (proximal tubule, distal convoluted tubule, and inner-medullary collecting duct) and basolateral membranes (thick ascending limb, distal convoluted tubule, cortical and outer-medullary collecting duct), with its highest expression being at the basolateral membrane of the cortical thick ascending limb [26]. Hence, we represent *v* for (i) paracellular permeability, NKCC2 activity, and renal outer-medullary potassium channel (ROMK) in the thick ascending limb, (ii) Na^+^-Cl^-^ cotransporter (NCC) activity in the distal convoluted tubule, (iii) H^+^-ATPase flux in outer-medullary collecting duct type A cells, and (iv) water permeability in the inner-medullary collecting duct. The parameters α_*v*,Ca_ and α_*v*,Mg_ are negative if CaSR has an inhibitory effect on *v* and positive otherwise. Since the effect of Mg^2+^ on CaSR activation is ∼50-66% of that of Ca^2+^, we set α_*v*,Mg_= 0.6α_*v*,Ca_. The value for α_*v*,Ca_ and α_*v*,Mg_ for each of the segments are given in Table 1.

## 3. Results

### 3.1 Baseline results

Figure X shows the segmental delivery, transport, and luminal fluid concentration of Mg^2+^ and Ca^2+^ in male rats. Results for other electrolytes and water can be found in Ref [27]. The majority of the filtered Ca^2+^ is reabsorbed along the proximal tubule, accounting for 68% of the filtered load; by contrast, Mg^2+^ reabsorption along this segment accounts for only 22% of the filtered load. Since Mg^2+^ reabsorption is low along the proximal tubule, Mg^2+^ concentration increases by ∼3-folds [4]. The majority of the overall Mg^2+^ transport occurs downstream along the cortical thick ascending limb (69% of the filtered load), where the lumen-positive membrane potential drives Mg^2+^ reabsorption via the paracellular pathway. The fractional reabsorption of Ca^2+^ along the medullary and cortical thick ascending limb is 20%. The final nephron segment that transports Mg^2+^ is the distal convoluted tubule, where 6.6% of the filtered Mg^2+^ is reabsorbed. The fractional reabsorption of Ca^2+^ along the distal convoluted tubule and connecting tubule is 7.9%. Finally, fractional urinary Mg^2+^ and Ca^2+^ excretions are 3.9% and 2.8%, respectively.

### 3.2 Effect of loop diuretics

Loop diuretics inhibit NKCC2, which is expressed on the apical membrane of the thick ascending limb. We simulated the effect of acute administration of loop diuretics by inhibiting NKCC2 activity by 70%. We assumed that the NKCC2 inhibitor was administrated for long enough to significantly impair the kidney’s ability to generate an axial osmolality gradient. The cortical interstitial concentrations were assumed to remain unchanged. Since the concentrating mechanism of the outer medulla is significantly impaired following complete NKCC2 inhibition, the interstitial concentration of Mg^2+^ at the outer-inner medullary boundary is lowered to 0.77 mM (from baseline value of ??mM). At the papillary tip, the interstitial concentration of Mg^2+^ is reduced to 0.48 mM. For changes in the interstitial concentrations of Na^+^, K^+^, Cl^-^, urea, and Ca^2+^ refer to [11,28].

The predicted Mg^2+^ and Ca^2+^ transport along the thick ascending limb and distal tubules and urinary Mg^2+^ and Ca^2+^ excretions following NKCC2 inhibition in male rats are shown in Fig. 3. Administration of furosemide to male rats increased Mg^2+^ excretion to 240% of control excretion value [29]. NKCC2 inhibition decreased fractional Mg^2+^ reabsorption along the cortical thick ascending limb to 57% from the baseline fractional reabsorption of 69% (Fig. 3). In male rats, this 12% decrease in Mg^2+^ reabsorption along the cortical thick ascending limb should ideally increase Mg^2+^ excretion to ∼430% of the control value. However, according to the experimental study [29], Mg^2+^ excretion increases to 240% of the control value following furosemide treatment. This indicates that there must be compensatory increase in Mg^2+^ reabsorption along the distal convoluted tubule. Indeed, TRPM6 mRNA expression was found to be increased by 30% in male mice undergoing furosemide treatment [30]. Our model simulations indicated that TRPM6 activity must be increased by 68% to account for the experimental increase in Mg^2+^ excretion. Fractional Ca^2+^ reabsorption along the thick ascending limb decreased by 17% following NKCC2 inhibition (Fig. 3). This resulted in urinary Ca^2+^ excretion increasing to 236% of the baseline excretion value (Fig. 2).

**Figure 1.**
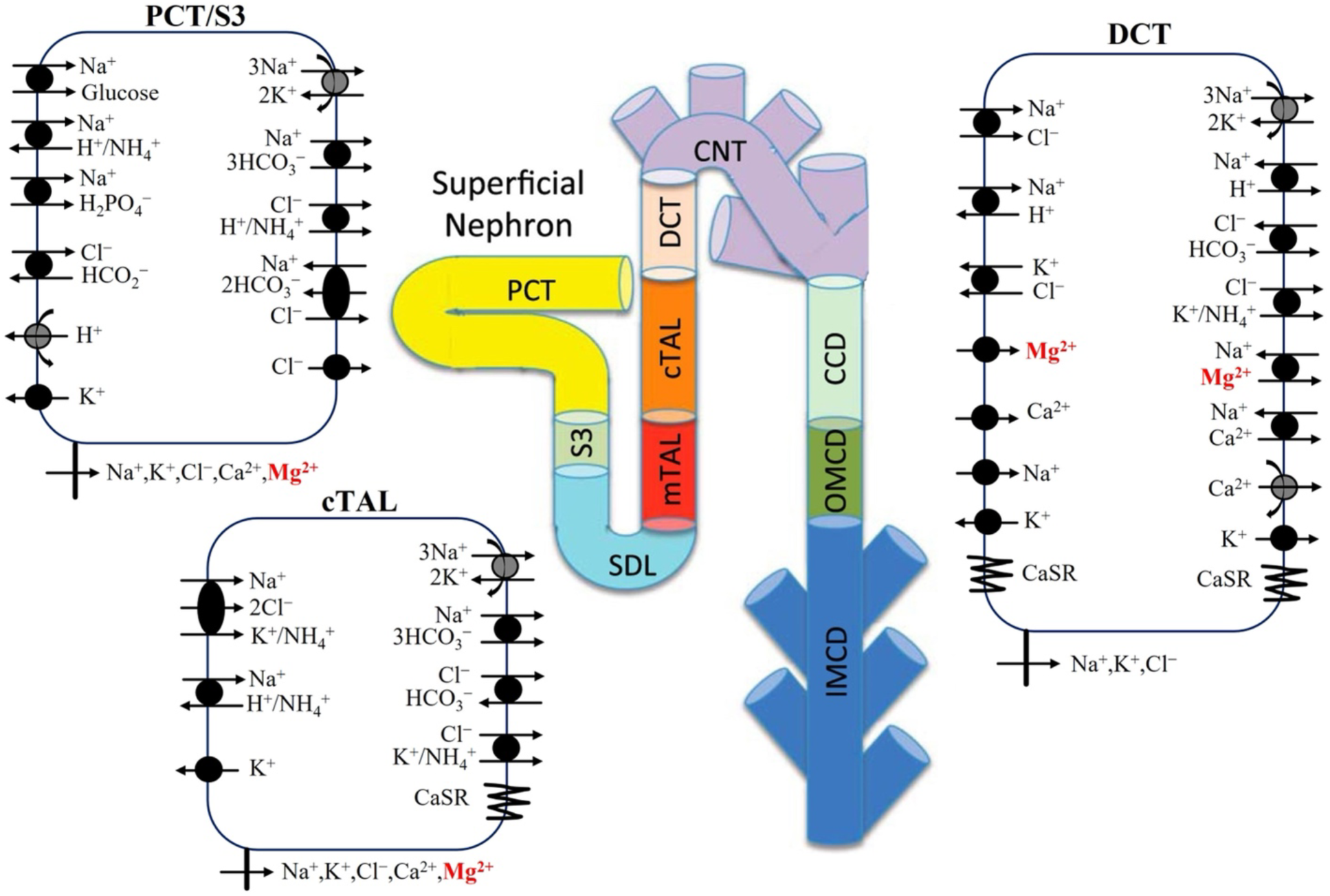
Model diagram of epithelial transport of Mg^2+^ and selected electrolytes along the superficial nephron. Mg^2+^ transport occurs along the proximal tubule (proximal convoluted tubule, PCT, and S3), cortical thick ascending limb (cTAL), and distal convoluted tubule (DCT). Only the major Na^+^, K^+^, Cl^-^, Ca^2+^, and Mg^2+^ transporters are shown. PCT, proximal convoluted tubule; SDL, short descending limb; mTAL, medullary thick ascending limb; limb; CNT, connecting tubule; CCD, cortical collecting duct; OMCD, outer-medullary collecting duct; IMCD, inner-medullary collecting duct.

**Figure 2:**
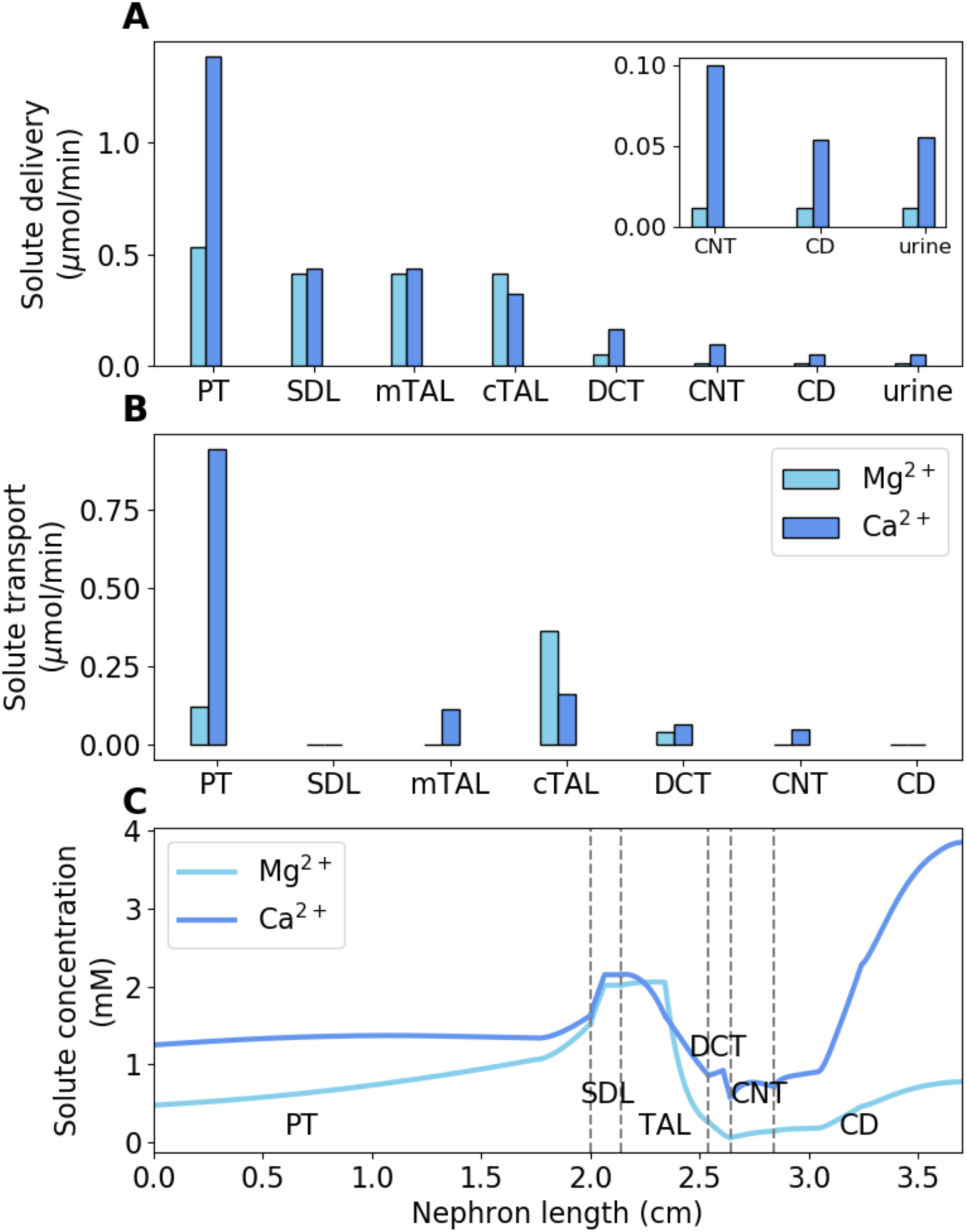
(A) Delivery of Mg^2+^ and Ca^2+^ to key nephron segments in male rats, given per kidney. (B) Mg^2+^ and Ca^2+^ transepithelial transport along key nephron segments in male rats, given per kidney. (C) Luminal Mg^2+^ and Ca^2+^ concentrations along key nephron segments in male rats. PT, proximal tubule; SDL, short descending limb; mTAL/cTAL, medullary/cortical thick ascending limb; DCT, distal convoluted tubule; CNT, connecting tubule; CD, collecting duct.

### 3.3 Effect of thiazide diuretics

Thiazide diuretics inhibit NCC, which is expressed along the apical membrane of the distal convoluted tubule. We simulated the effect of acute administration of thiazide diuretics by inhibiting NCC activity by 70%. In the NCC inhibition simulations, baseline interstitial concentration profiles were used.

Administration of bendrofluazide, a thiazide diuretic, did not cause any significant change in Mg^2+^ excretion in male rats [31]. In agreement to the experimental data, the predicted fractional Mg^2+^ excretion after NCC inhibition increased to 3.1% from the baseline value of 2.8% (Fig. 3). Acute administration of chlorothiazide to male mice increased TRPV5 mRNA expression by 40-80% and decreased Ca^2+^ excretion by ∼60% [32]. Accordingly, we increased TRPV5 activity by 52% following NCC inhibition in our model. This resulted in the predicted Ca^2+^ excretion to decrease by 63% from the baseline excretion value (Fig. 2).

**Figure 3:**
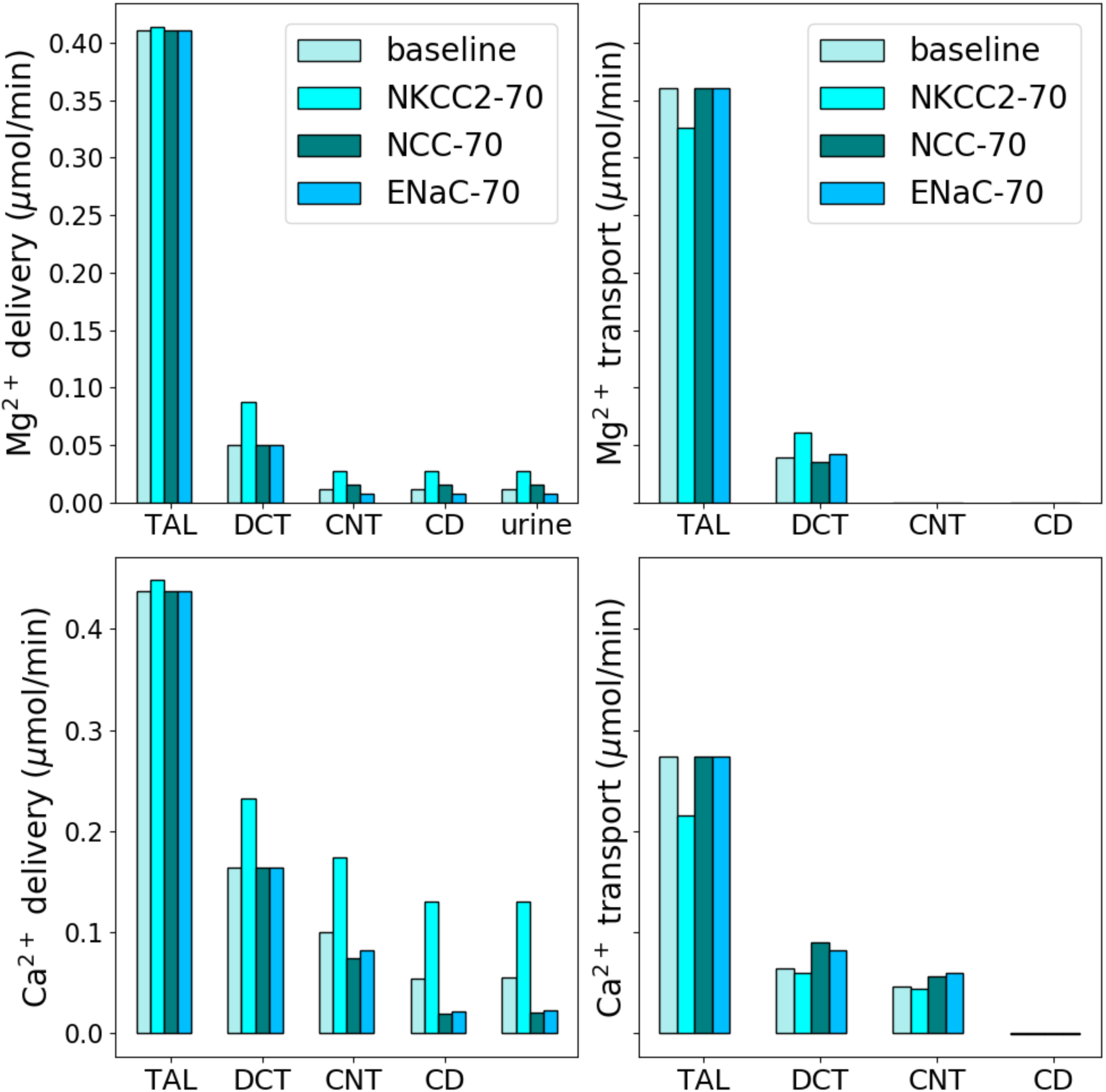
Delivery and transport of Mg^2+^ and Ca^2+^ along key nephron segments in male rats under normal condition and 70% inhibition of NKCC2, NCC, and ENaC. The values are given per kidney. Notations are analogous to Fig. 1.

### 3.4 Effect of K-sparing diuretics

K-sparing diuretics, such as amiloride, block Na^+^ uptake through ENaC, expressed on the apical membrane of the late distal convoluted tubule as well as along the full length of the connecting tubule and collecting ducts. K-sparing diuretics are weak diuretics often used in combination with other diuretics to lower blood pressure. By inhibiting ENaC, these diuretics hyperpolarize the luminal membrane potential and increase K^+^, Ca^2+^, and Mg^2+^ uptake [33–36]. In our model, we simulated the effect of K-sparing diuretics by reducing ENaC activity by 70%.

K-sparing diuretics, amiloride and triamterene, reduced Mg^2+^ excretion in male rats by ∼40% [31]. By inhibiting ENaC, these diuretics hyperpolarize the luminal membrane potential and increase K^+^, Ca^2+^, and Mg^2+^ uptake [33–36]. Our model simulations predicted Mg^2+^ reabsorption along the distal convoluted tubule to increase by 8.2% and Ca^2+^ reabsorption along the distal convoluted tubule and connecting tubule to increase by 29% (Fig. 3). These increased reabsorptions decreased Mg^2+^ and Ca^2+^ excretions by 31% and 56%, respectively (Fig. 3).

## 4. Discussion

Calcium (Ca^2+^) and magnesium (Mg^2+^) are both essential for cellular function. The homeostasis of these cations must be tightly regulated, and that balance is facilitated by intestinal absorption and renal excretion. For Na^+^, Cl^-^, K^+^, Ca^2+^, and many other major filtered solutes, most of the renal reabsorption occurs along the proximal convoluted tubule (about 1/2 to 2/3 in rats); the same is true for water. Thus, the luminal concentrations of these solutes, including Ca^2+^, remain close to plasma along the proximal convoluted tubule. The majority of proximal tubule Ca^2+^ reabsorption occurs via a passive paracellular process, driven by Na^+^ reabsorption mediated primarily by NHE3, and by subsequent water reabsorption. In contrast, only 15-25% of the filtered Mg^2+^ load is reabsorbed along the proximal tubule. As a result, its concentration rises significantly along the proximal tubule. For Mg^2+^, most of the reabsorption occurs along the *cortical* thick ascending limb (about 60-70%), while somewhat unexpected, essentially none along the medullary thick ascending limb. Most of the remainder of the Mg^2+^ is reabsorbed along the distal convoluted tubule.

What difference does it make for the cortical thick ascending limb and distal convoluted tubule to handle most of the Mg^2+^ transport, instead of the proximal tubule, as in the case of Na^+^ and Cl^-^? Having these distal segments responsible for transporting a substantial fraction of the filtered Mg^2+^ load via the pathways that can be regulated may give the kidney a better ability to regulate Mg^2+^ balance. Recall that plasma Mg^2+^ level is orders of magnitude lower than Na^+^ or Cl^-^. Thus, to maintain plasma [Mg^2+^] within a narrow range, the ability to fine tune renal Mg^2+^ transport is particularly crucial. Parathyroid hormone, for instance, increases Mg^2+^ reabsorption in both the cortical thick ascending limb and distal convoluted tubule [37]. Transport of Mg^2+^ along these segments can also be regulated by hormones such as calcitonin, vasopressin, glucagon, and β-adrenergic agonists [38]. Coincidentally, some common diuretics also target these segments.

A goal of this study is to better understand the impact of diuretics on renal Ca^2+^ and Mg^2+^ transport. Diuretics are medications that reduce fluid buildup in the body and are often employed in the management of hypertension, edema, and various other conditions influenced by changes in electrolyte transport. In the context of kidney function, diuretics often focus on transport proteins or mechanisms vital for the reabsorption of Na^+^, Cl^-^, and water. Considering that renal Mg^2+^ transport is driven primarily by the electrochemical gradients established by tubular NaCl transport processes, potential modifications in the renal handling of Ca^2+^ and Mg^2+^ by the administration of diuretics deserve scrutiny.

Loop diuretics such as furosemide induce notable natriuresis by targeting the thick ascending limb of the nephron; specifically, they inhibit NKCC2-dependent transport by competing for the chloride (Cl^-^) binding site [39]. Given that the inhibition of NKCC2 reduces the lumen-positive transepithelial voltage gradient across the thick ascending limb epithelium [40], it is unsurprising that the transport of magnesium ions (Mg^2+^) in this segment is substantially administration [41,42]. In fact, loop diuretics like furosemide induce hypercalciuria and hypermagnesiuria in both experimental animals [43] and human subjects [44]. Thiazide diuretics indue a natriuretic response by inhibiting NCC and blocking NaCl transport in the distal convoluted tubule. A hypocalciuric effect has been reported following thiazide treatment [45]. Consistent with the drug’s mode of action, among hypertensive individuals undergoing chronic thiazide treatment, there is a slight decrease in serum Mg^2+^ levels compared to those not taking diuretics [46], although the effect appears to be subtle. K^+^-sparing diuretics function by inhibiting ENaC, a protein expressed in the late distal convoluted tubule, connecting tubule, and collecting duct of the kidney. Research has confirmed that K^+^-sparing diuretics impact the urinary excretion of Ca^2+^ and Mg^2+^ in both human subjects and animals. Hypertensive individuals undergoing K^+^-sparing diuretics treatment exhibit reduced urinary Ca^2+^ excretion [46] and elevated serum Mg^2+^ levels compared to those not receiving treatment [46]. Model simulations of the administration of these diuretics predict marked diuretic and natriuretic effects, with the strongest response induced by loop diuretics (Fig. 3).

The present study considers renal Ca^2+^ and Mg^2+^ transport under normal physiological conditions. The homeostasis of these electrolytes is altered pregnancy, lactation, and dietary restriction, as well as in diseases such as diabetes and chronic kidney disease. To utilize the present model for *in silico* studies of how the kidney adapts in terms of Ca^2+^ and Mg^2+^ transport under these physiological and pathophysiological conditions, one can combine the model with computational models of kidney function for a pregnant rat [47,48], a diabetic rat [49–51], a nephrectomized rat [9,52], or a whole-kidney model [53–55]. The resulting models may provide insights into altered renal Ca^2+^ and Mg^2+^ transport under these conditions. How do changes in renal Mg^2+^ transport impact whole-body Mg^2+^ homeostasis? A whole-body Mg^2+^ balance model may help answer that question. By incorporating the present model into whole-body electrolyte balance models (e.g., [56–59]), one can obtain an integrative model to study whole-body Mg^2+^ balance.

## Code availability

The code used for this study can be accessed at https://github.com/Pritha17/Nephron-Mg_Ca_transport.

## Acknowledgements

This work was supported by the Canada 150 Research Chair program, National Sciences and Engineering Research Council of Canada (NSERC) Discovery Grant, and Canada Institutes of Health Research (CIHR) Project Grant (to A.T.L).

## Author contributions

Conceptualization, A.T.L.; Methodology, P.D. and A.T.L.; Software, Validation, Formal Analysis, and Investigation, P.D.; Resources, A.T.L.; Data Curation, P.D.; Writing – Original Draft, P.D. and A.T.L; Writing – Review & Editing, P.D., S.H., and A.T.L.; Visualization, P.D.; Supervision, P.D. and A.T.L.; Funding Acquisition, A.T.L.

